# Systemic delivery of a splice-switching oligonucleotide heteroduplex corrects splicing in central nervous system and muscle in spinal muscular atrophy mice

**DOI:** 10.1101/2024.01.24.577012

**Authors:** François Halloy, Nina Ahlskog, Matthew Wood

## Abstract

Oligonucleotide therapeutics are an established class of drugs for the treatment of genetic disorders. Their clinical development is challenging, however, as they typically distribute poorly to extra-hepatic tissues after systemic injection. Here we tested the heteroduplex oligonucleotide (HDO) platform for systemic delivery of *SMN2* splice-switching oligonucleotides of 2’-*O-*methoxyethyl phosphorothioate or phosphorodiamidate morpholino oligomer chemistries. We first showed that splice-switching HDO cargoes correct *SMN2* splicing in cells derived from spinal muscular atrophy (SMA) patients, and validated extra-hepatic activity in spinal cord and muscle in a mouse model of SMA following systemic delivery. Our study raises prospects for delivery of nusinersen, the 2’-*O-*methoxylethyl phosphorothioate oligonucleotide therapy approved for SMA and currently delivered by intrathecal injection, by systemic injection exploiting the HDO chemistry platform. Our findings also suggest that oligonucleotide drugs lacking convincing *in vivo* efficacy in muscle tissue could be delivered effectively by the HDO technology.

## INTRODUCTION

Oligonucleotides (ONs) are a major class of drugs in human pharmacopeia, with 16 ONs approved by regulatory authorities to date for the treatment of various genetic disorders (1). ONs are defined as short, synthetic nucleic acids with an average length of 18-30 nucleotides (2). In the cell, ONs bind a cognate sequence of RNA in the cell according to Watson-Crick base-pairing rules, whereupon they modulate gene expression (3). ONs can be broken down into three main categories (4). RNase-H recruiting oligonucleotides, also known as gapmers, are single-stranded ONs that bind a target RNA transcript, causing its cleavage by RNase H1. A second class, called small interfering RNAs (siRNAs), encompasses duplexes of an antisense and a sense strand. siRNAs are bound by the RISC complex *in cellulo*, whereupon the duplex is dissociated and the active strand binds a target transcript, ultimately causing target cleavage (5). The third group, labelled as “steric-blocking ONs”, binds a target transcript without cleavage activity but rather prevents its interaction with RNA-binding proteins or other cognate RNA species (3). Steric-blocking ONs encompass anti-microRNAs, which bind non-coding transcripts named microRNAs (6), and splice-switching oligonucleotides (SSOs), which bind pre-messenger RNA transcripts (7).

SSOs are of particular interest for genetic diseases caused by mutations altering RNA splicing. Indeed, these mutations lead to the disruption or creation of splice sites, which impacts mRNA stability or the functionality of proteins. Among the 16 approved ON drugs, five are SSOs: nusinersen is approved for the treatment of spinal muscular atrophy (SMA) (8), while casimersen, golodirsen, viltolarsen and eteplirsen are approved for the treatment of Duchenne Muscular Dystrophy (DMD) (9-12). Nusinersen is fully modified with 2’-*O*-methoxylethyl (MOE) riboses and a phosphorothioated backbone, whereas casimersen, golodirsen, viltolarsen and eteplirsen are phosphorodiamidate morpholino oligomers (PMOs). The MOE and PMO modifications are manmade chemistries conferring superior stability *in vivo*, typically in weeks, and high binding affinity compared to unmodified nucleic acids.

SMA is a genetic disorder of the CNS and peripheral nervous system that affects the viability of motor neurons due to loss-of-function mutation in the Survival Motor Neuron 1 (*SMN1*) gene (13). Humans have a paralog gene, *SMN2*, that exists in various copy numbers (14).

Due to exon-skipping in the *SMN2* pre-mRNA, the primary product of *SMN2* mRNA lacks exon 7 (*SMN2-Δ7*) and is unstable and rapidly degraded (15). As only ≈10% of the *SMN2* mRNA is full length and translated to functional SMN protein, the copy numbers are inversely related to phenotype severity in SMA patients (16). Increasing the translation efficiency of *SMN2* through SSOs is a validated therapeutic approach that led to the development of nusinersen.

In spite of their growing success, important challenges remain in ON development, such as delivery into disease-relevant extra-hepatic tissues (17). The delivery issue stems from the intrinsic chemical properties of ONs, which are polar, large, and polyanionic macromolecules at physiological pH. An established route for delivery of ON therapies is systemic injection, e.g. intravenously (i.v.) or subcutaneously (s.c.). Systemic injections are relatively low-risk and enable rapid ON biodistribution into tissues through the bloodstream (18,19). Many tissues, however, are refractory to ON uptake, with muscle (19) and brain (2) being notorious examples. While delivery into central nervous system (CNS) can be achieved by local injection, *e*.*g*., intrathecal route for approved drugs nusinersen or tofersen (20), therapies for muscle indications rely on systemic injections, with suboptimal efficacy (9). Overall, the lack of uptake and ultimately efficacy in muscle and CNS tissues has caused many ON candidates to fail in pre-clinical and clinical programs (17).

Improving ON delivery into tissues is a continuing concern within the field. An established strategy is ON conjugation to moieties enhancing cellular uptake. Fatty acids and lipids (21), peptides or antibodies (22) have been widely used for this purpose. Particularly successful examples are N-acetylgalactosamine (GalNAc) conjugates. GalNAc conjugates enhance ON delivery into hepatocytes in the liver by binding a receptor specifically expressed on these cells, the asialoglycoprotein receptor. Several GalNAc-ON conjugates are now approved in clinics for liver disorders (23). Other significant delivery approaches rely on modifying ON backbone (24) or encapsulating ONs (25).

In a seminal 2015 study, the Yokota group introduced the heteroduplex oligonucleotide (HDO) platform as a new means to improve ON delivery (26). HDOs are made of an active ON strand, e.g., a gapmer-, a splice-switching-or an antimiR oligonucleotide, and a fully complementary carrier strand modified at the 5’ end with a lipid. The HDO scaffold can be seen as a pro-drug enhancing ON delivery, and activity, *in vivo* (27). In another 2021 study, the Yokota group showed that gapmer HDOs targeting the non-coding RNA *Malat1* crossed the blood-brain-barrier after systemic injection and knocked-down *Malat1* in the CNS of monkeys and rodents (28). Knock-down was achieved at doses of 25 to 50 mg active strand/kg animal body weight which, interestingly, is in the range of concentrations used in SSO studies (29,30). We therefore hypothesised that we could use the HDO scaffold to deliver *SMN2* SSOs for SMA therapy. This approach would have several advantages.

Nusinersen is currently delivered by intrathecal injection, directly into the CNS compartment and repeated intrathecal injections can be challenging in SMA patients who often present a deformed spine (31). They are also riskier than systemic injections, with infection, nerve injury and leakage of cerebrospinal fluid all potential issues (32). Delivering nusinersen preparations through an HDO scaffold and systemic injections would alleviate these risks.

Furthermore, *SMN2* HDOs delivered by systemic injections could reach peripheral tissues in addition to the CNS, which was shown by Krainer and colleagues to be essential for long-term rescue of SMA (29).

Here we tested HDO constructs of PMO and MOE-PS chemistry for *SMN2* splice correction. We verified HDO activity *in vitro* in patient-derived cells and elected the MOE-PS HDO for *in vivo* validation in an humanized adult mouse model of SMA. Our results show that a single intravenous injection of MOE-PS HDO at 40 mg/kg increased full-length *SMN2* transcript levels by ∼20% in spinal cord and ∼100% in cardiac and skeletal muscle. Our findings pave the way for further development of HDO therapeutics for SMA and other neuromuscular disorders.

## RESULTS

### Splice-switching HDOs correct *SMN2* splicing in SMA Type II patient fibroblasts

The first step of this study was to design heteroduplex oligonucleotides for *SMN2* splice correction. We selected a 25-nt PMO sequence, named **PMO** (5’ GTAAGATTCACTTTCATAATGCTGG 3’), for which we previously validated activity on *SMN2* splicing and exon 7 inclusion *in vivo* in mice (33). **PMO** is slightly longer than nusinersen, the 18-nt approved drug for SMA (8), but binds the same locus and encompasses the nusinersen binding site. We designed the carrier strand **Cs** to **PMO** as to match closely the design published by Nagata and colleagues (28). **Cs** was fully complementary to PMO, had a central stretch of phosphodiester RNA, phosphorothioated 2’-O-methyl wings, and a 5’-triethylene glycol-cholesteryl modifier (**Figure 1A**). We were also interested in comparing the efficacies of PMO and *2’-*O-MOE chemistries for *SMN2* splice correction, and designed **MOE**, a 2’-*O-*MOE phosphorothioate version of **PMO** (**Figure 1B**). For both chemistries, we performed duplex annealing in RNAse-free water and confirmed formation of **MOE**HDO and **PMO**_HDO_ by polyacrylamide gel electrophoresis (PAGE) on a Tris-Borate-EDTA native gel (**Figures 1C** and **1D**, respectively).

**Figure 1.**
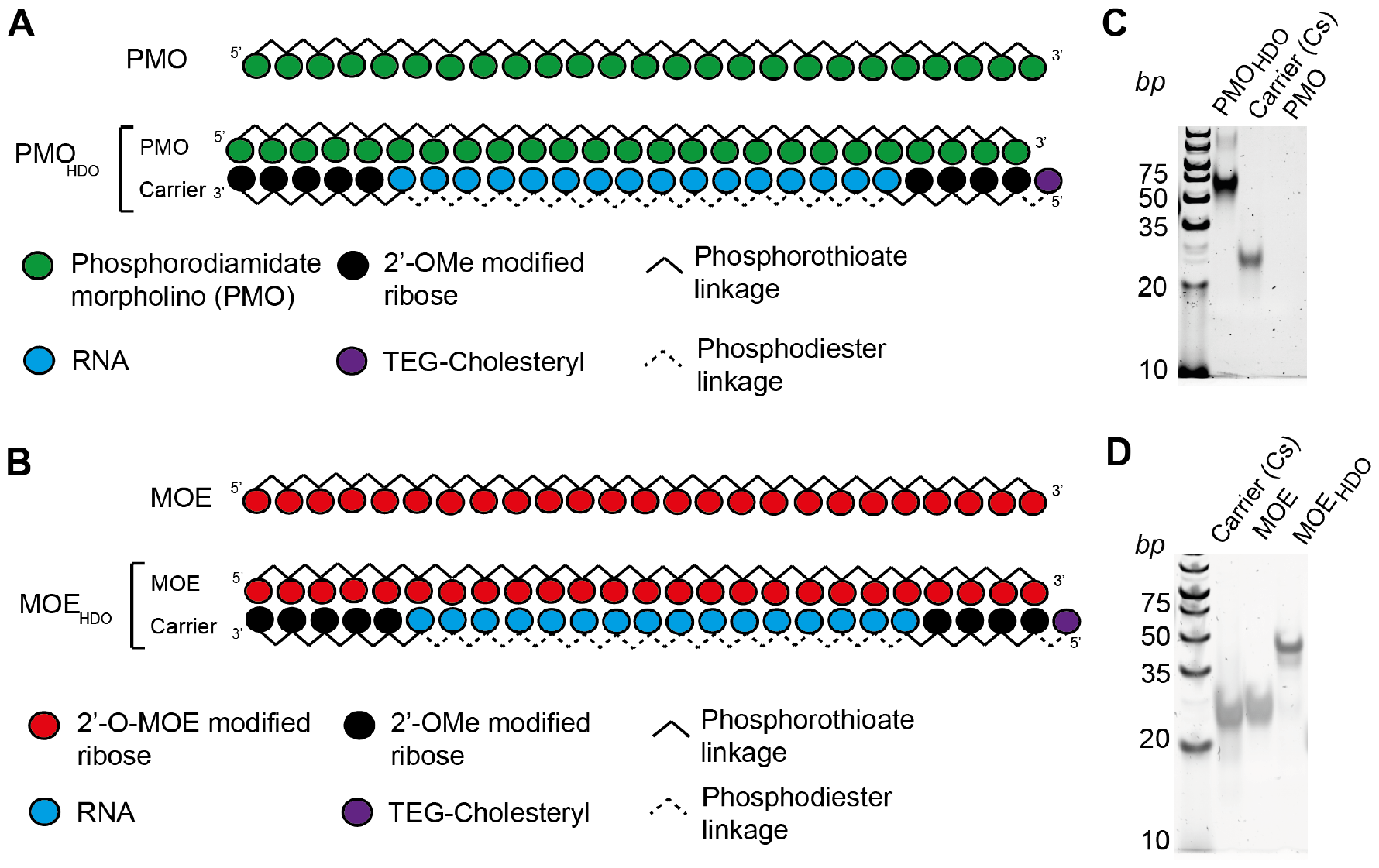
Design of splice-switching heteroduplexes for SMN2 splice correction. We designed 25-nucleotide HDO versions of a known PMO oligonucleotide, **PMO** (**A**) and of its 2’-O-MOE phosphorothioate version **MOE** (**B**) for *SMN2* splice correction. The carrier strand **Cs** was modified with a 5’-TEG cholesteryl moiety, 2’-*O*-Me phosphorothioated wings, and a central RNA phosphodiester core. We confirmed annealing by PAGE gel electrophoresis in native conditions on a 15% Tris-Borate-EDTA gel for both **PMO**_HDO_ (**C**) and **MOE**_HDO_ (**D**). TEG: triethylene glycol; 2’-O-MOE: 2’*-O*-methoxyethyl ribose; 2’-*O*-Me: 2’*-O-*methyl ribose.

We then turned to testing of **PMO**_HDO_ and **MOE**_HDO_ in an SMA Type II fibroblast cell line enabling splice correction on endogenous *SMN2* transcripts. Activity was assessed 24 h post treatment by calculating relative expression of full-length *SMN2* normalised to total *SMN2* expression in quantitative reverse transcription polymerase chain reaction (RT-qPCR). We tested **PMO, PMO**_HDO_ and a negative control **PMO**_**c**trl_ at concentrations of 50 nM, 500 nM and 1 µM, based on a previous account (34). We observed a dose-dependent activity for both **PMO** and **PMO**_HDO_ (**Figure 2A**). **PMO** significantly increased *SMN2* transcripts levels at 1 µM concentration, with a maximal 1.56-fold increase **PMO**_HDO_ was similarly active, with 1.65- and 1.58-fold increases attained at 500 nM and 1 µM concentration, respectively. This pattern was confirmed by calculating relative expression of full-length *SMN2* transcripts and using *ACTB* as internal control (**Figure S1A**). There, we obtained 1.64- and 1.67-fold increases in full length *SMN2* ratios at 1 µM concentration for **PMO** and **PMO**_HDO_, respectively.

**Figure 2.**
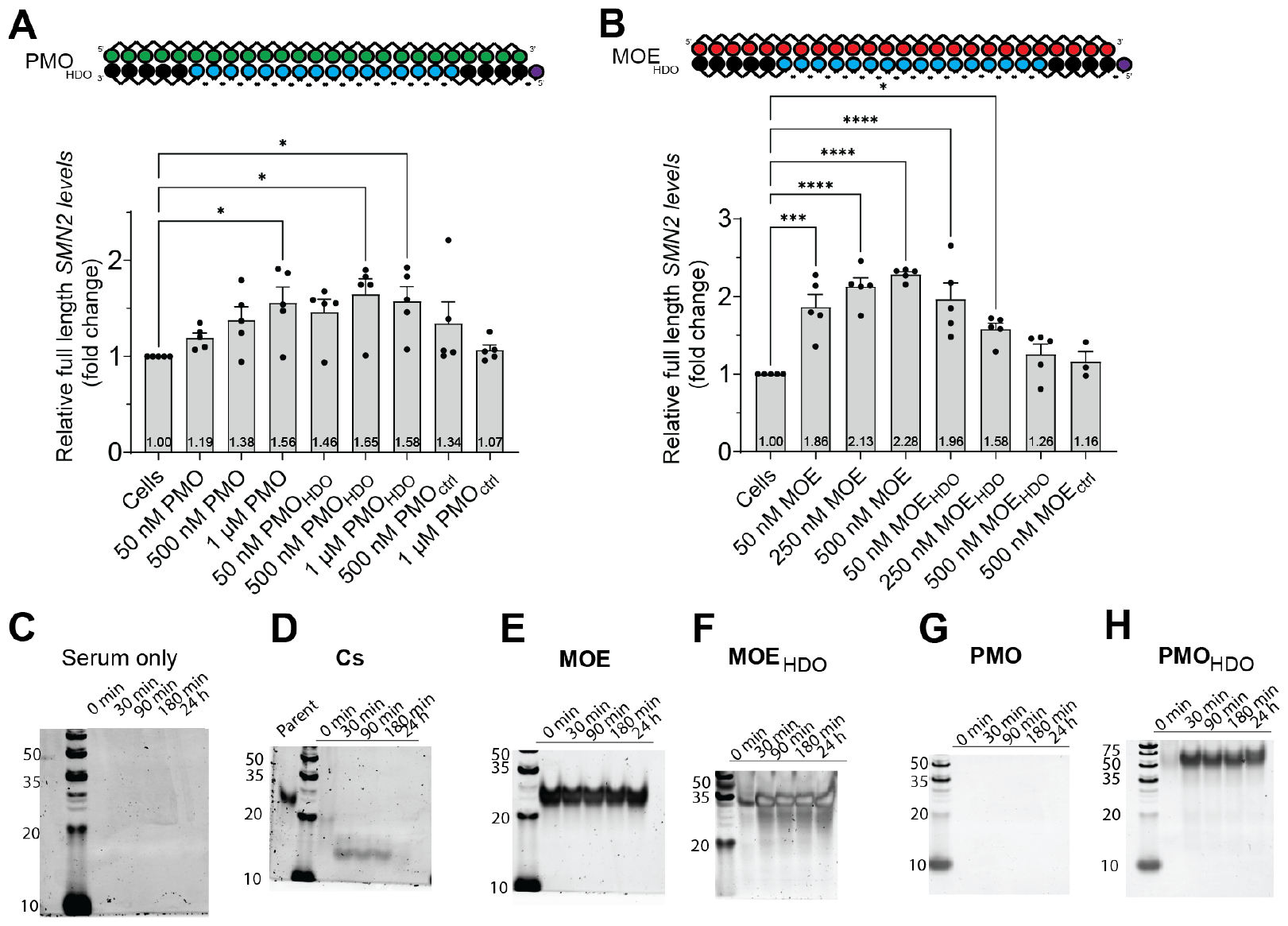
In vitro activity of SMN2 splice-switching HDOs and stability in serum. **(A-B)** Oligonucleotide activity was assessed in SMA Type II fibroblast cells after 24h treatment at indicated concentrations, by means of RT-qPCR assays detecting full-length *SMN2* and total *SMN2* transcripts. Expression levels of full length *SMN2* were normalised to total *SMN2* transcripts. **A** Cells were treated with increasing concentrations of **PMO, PMO**_HDO_ or **PMO**_ctrl._ *(n)* = 5 biological replicates; data expressed as mean value (SEM: standard error of the mean); statistical analysis was conducted by comparison to the untreated group with one-way Anova (Dunnett’s correction). **B** Cells were treated with increasing concentrations of **MOE, MOE**_HDO_ or **MOE**_ctrl_ *(n)* = 3-5 biological replicates; data expressed as mean value (SEM: standard error of measurement); statistical analysis was conducted by comparison to the untreated group with one-way Anova (Dunnett’s correction). **(C-H)** Oligonucleotides were incubated in mouse serum at indicated time points and analysed by PAGE after proteinase K digestion. **C** no oligonucleotide-derived bands were obtained in the serum control; **D** degradation in shorter fragments was observed for **Cs**; **E-F**: no degradation was observed for **MOE** and **MOE**_**HDO**_; **G** no band could be detected for **PMO**, which we hypothesised was due to lack of staining by SYBR Gold; **H**: no degradation was observed for **PMO**_**HDO**._

**MOE**_HDO_ was tested against the parent control **MOE** and a six-mismatch negative control **MOE**_ctrl_ (35) at increasing concentrations of 50, 250 and 500 nM. (**Figure 2B**). **MOE** was active at all three concentrations and exhibited a dose-dependent response with a maximum 2.31-fold increase in full-length *SMN2* transcripts at the 500 nM dose. We observed statistically-significant activity for **MOE**_HDO_ treatments, with a maximal ∼ 2-fold increase in full-length *SMN2* transcripts at the lowest dose (50 nM) and 1.58-fold increase at 250 nM. Findings were broadly similar when the RT-qPCR data were normalised to expression of a endogenous reference gene (*ACTB*) (**Figure S1B**). There, **MOE** elicited a maximal 3.3-fold increase in full-length *SMN2* at 500 nM concentration. Small increases in full length *SMN2* were observed for treatment with **MOE**_HDO_ (1.41-fold at 50 nM concentration, and 1.34-fold at 250 nM concentration), although this did not reach statistical significance at the p < 0.05 level.

**MOE**_HDO_ and **PMO**_HDO_ were also non-toxic at the concentrations tested, according to a commercial viability assay (**Figure S2**). Overall, these results indicated that both **MOE**_HDO_ and **PMO**_HDO_ were functional in cells and elicited a biological response on *SMN2* transcript levels. **PMO** and **PMO**_HDO_ appeared similar in activity, whereas **MOE** and **MOE**_HDO_ showed opposite dose-response patterns, with **MOE**_HDO_ most active at the two lowest concentrations.

### *In vitro* stability of PMO_HDO_ and MOE_HDO_ in mouse serum

It is well established from a number of studies that 2’-*O-*MOE PS and PMO chemistries are resistant to nuclease degradation for days in plasma or in tissues (36-38). **Cs**, however, encompasses a 16-nt stretch of unmodified RNA whose stability toward nucleases is limited. Hence, in anticipation of *in vivo* studies, we set out to determine if **MOE**_HDO_ and **PMO**_HDO_ were stable in mouse serum for up to 24h. Single strands and duplexes were spiked in mouse serum and stability was assessed by polyacrylamide gel electrophoresis after 30 min, 90 min, 180 min, or 24h incubation (**Figure 2**, panels **C-H**). We first verified that no bands were detected in mouse serum in the 10-50 bp range (**Figure 2C**). As anticipated, **Cs** alone was rapidly degraded in shorter fragments, and no clear band could be resolved by the last 24h timepoint (**Figure 2D**). This result appears consistent with initial cleavage of the central phosphodiester RNA core by endonucleases, followed by slower cleavage of the 2’-O-Me wings. **MOE** and **MOE**_HDO_ appeared undegraded under the conditions and timepoints tested (**Figure 2E** and **2F**). No band was detected for **PMO**, which we assumed was due to a lack of staining of single-stranded PMO compounds by SYBR Gold dye (**Figure 2G**). However, we confirmed stability of **PMO**_HDO,_ with no metabolite species arising from **Cs** observed up to 24h incubation (**Figure 2H**). Taken together, our results suggested that **MOE**_HDO_ and **PMO**_HDO_ would remain sufficiently stable in the bloodstream to have an impact on biodistribution.

### MOE_HDO_ corrects splicing and *SMN2* expression in vivo in adult SMA mice

Based on their superior results *in vitro* (**Figure 2B**), we selected **MOE** and **MOE**_HDO_ for *in vivo* investigation. To this end, we used a transgenic mouse model carrying four copies of the human *SMN2* gene. This model has low clinical severity and enables activity studies in adult animals, with a fully developed blood-brain-barrier (BBB). Previous research has indeed established that mouse BBB is fully matured at 12-24 days of age (39). More severe, and short-lived mouse models of SMA (40) require injections directly after birth and are therefore not good proxies for delivery beyond a mature BBB.

We opted for a single dose of **MOE** or **MOE**_HDO_ at 40 mg active strand/kg intravenously in the tail vein. No side effects were observed with **MOE**. Transient side effects, *i*.*e*. spasms and lethargy, were observed upon injection of **MOE**_**HDO**_ and fully subsided within 30-90 minutes. One mouse had incomplete recovery two hours post injection, however, and was culled according to our licence guidelines. We assessed activity and clinical chemistry biomarkers seven days post injection (34). We found no significant change in animals in a panel of kidney and liver toxicity biomarkers **(Figure S3)**.

We monitored *SMN2* splice correction levels in selected tissues using the RT-qPCR assay described above as well as a semi-quantitative PCR assay for detection of full-length and truncated (Δ7) *SMN2* transcripts. We first investigated splice correction in CNS tissues (**Figure 3**). In the brain, we did not detect activity in cerebellum, brain stem, or cortex.

**Figure 3.**
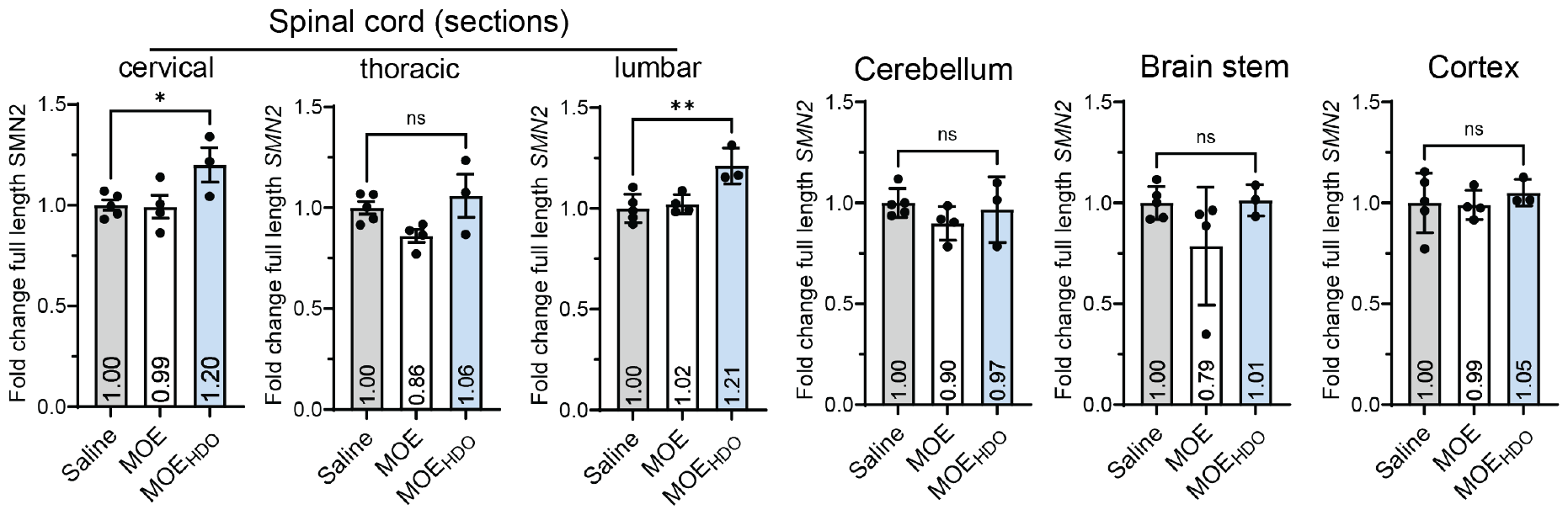
Central nervous system activity of MOE and MOE_HDO_ after intravenous injection at 40 mg/kg in adult mice carrying the human SMN2 gene. Activity was assayed with a RT-qPCR assay for quantification of correct *SMN2* transcripts and normalization to the reference transcript *PolJ*. Animal cohort size: (*n*)= 3-5 per group, one dot represents an independent animal; statistical significance was determined by comparison to the saline group using one-way ANOVA with Dunnett’s correction; * = *p* < 0.05, ** = *p*< 0.01, data expressed as mean value (SEM).

Pleasingly, however, **MOE**_HDO_ corrected *SMN2* splicing in spinal cord, with modest activity in cervical and lumbar sections at a 1.20- and 1.21-fold increase in full length *SMN2* transcripts, respectively. The effect was not observable in a semi-quantitative PCR assay for the visualisation of correct and Δ7 *SMN2* transcripts on agarose gel **(Figure S4)**, which is in part not surprising given the small effect size and the relative insensitivity of this methodology.

We then investigated activity in peripheral tissues (**Figure 4**). In heart, **MOE**_HDO_ selectively increased full-length *SMN2* transcript levels by a factor 1.56, while **MOE** showed no statistically significant effect (**Figure 4A**). **MOE**_HDO_ activity was further enhanced in skeletal muscle (gastrocnemius, tibialis anterior and quadriceps) with a consistent 2.1-2.2-fold increase. We only detected a moderate activity for **MOE** in tibialis anterior and quadriceps with a 1.6-fold increase and in gastrocnemius with a 1.4-fold increase. Supporting results were obtained in the semi-quantitative PCR assay detecting truncated and full-length *SMN2* transcripts (**Figure 4B**). There, a 30% increase in the ratio of correct *SMN2* transcripts was obtained with **MOE**_HDO_ in heart, and a ≈80% increase in tibialis anterior, gastrocnemius and quadriceps muscle. We finally assayed activity in kidney and liver (**Figure S5**). In the liver, **MOE**_HDO_ was significantly most active than **MOE**, whereas in kidney both compounds had similar levels of activity according to RT-qPCR and semi-quantitative PCR data (**Figure S5A** and **S5B**, respectively).

**Figure 4.**
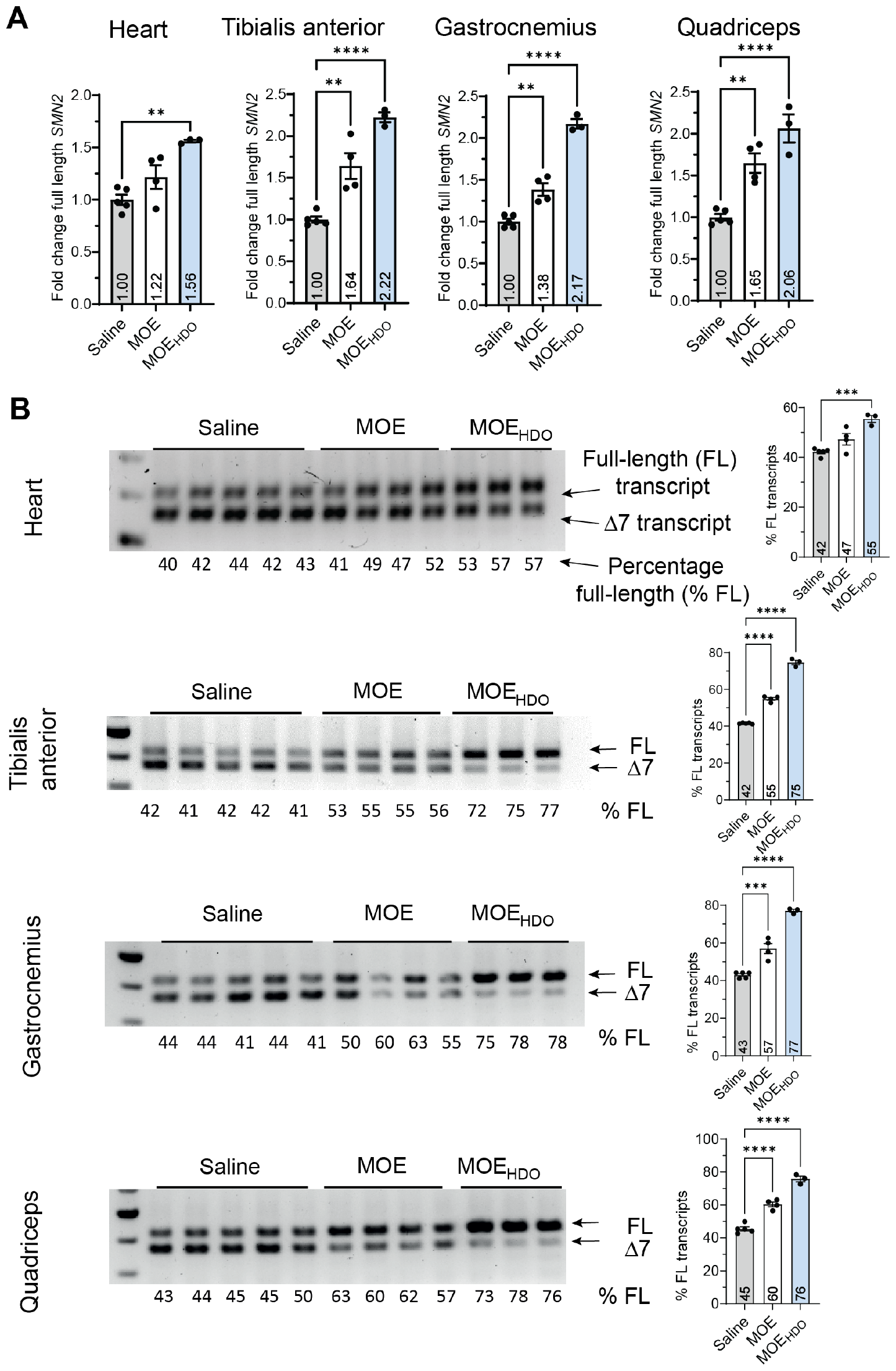
Muscle activity of MOE and MOE_HDO_ after intravenous injection at 40 mg/kg in adult mice carrying the human SMN2 gene. **A** Activity was assayed with a RT-qPCR assay for quantification of correct *SMN2* transcripts and normalization to the reference transcript *PolJ*, and **B** with a semi-quantitative PCR assay for simultaneous detection of full length and Δ7 *SMN2* transcripts. Animal cohort size was (*n*)= 3-5 per group, one dot represents an independent animal; statistical significance was determined by comparison to the saline group via one-way ANOVA with Dunnett’s correction; * = *p* < 0.05, ** = *p*< 0.01, *** = *p* < 0.001; **** = *p* < 0.0001; data expressed as mean value (SEM).

## DISCUSSION

Improving ON delivery in the context of neuromuscular disease is an active field of research (33,41,42). HDOs have emerged as a promising delivery vehicle in recent years (27), but no study to date had investigated HDOs in the context of SMA and *SMN2* splice correction.

Here, we tested whether HDOs carrying a splice-switching oligonucleotide of 2’-*O*-MOE PS chemistry (**MOE**_HDO_) or PMO chemistry (**PMO**_HDO_) were functional in diseased cells and selected **MOE**_HDO_ for *in vivo* investigation in a transgenic mouse model of SMA.

*In vitro*, our data showed that both **MOE**_HDO_ and **PMO**_HDO_ duplexes were biologically functional and that the active strand could successfully engage *SMN2* pre-mRNA. While **PMO** and **PMO**_HDO_ had comparable activities in SMA Type II fibroblasts in the conditions tested (**Figure 2A**), **MOE** and **MOE**_HDO_, exhibited opposite dose-response relationships (**Figure 2B**). We used assisted delivery with Lipofectamine 2000 for cell culture experiments, and it is well possible that these rankings in activity are due to distinct abilities of Lipofectamine reagent to deliver increasing concentrations of double-stranded structures with a fully polyanionic backbone (**MOE**_HDO_) or a partially uncharged backbone (**PMO**_HDO_).

Further studies are also needed to understand how **MOE**_HDO_ and **PMO**_HDO_ are processed in cells and how the active strand is released from the duplex prior to engaging its target. Thus far, previous studies with gapmer or antimir HDOs have demonstrated that either duplex unwinding or cleavage can occur (43,44), but this would need to be confirmed for splice-switching oligonucleotides.

We elected **MOE**_HDO_ for mouse testing as it elicited the best splice correction in realtime PCR (**Figure 2B**) and to stay as close as possible to the ON sequence and chemistry approved for SMA with nusinersen (8). **MOE** and **MOE**_HDO_ were injected intravenously and tissues harvested seven days post injection, following an established procedure (34). We found **MOE**_HDO_ modestly corrected *SMN2* splicing in spinal cord but not in brain itself (**Figure 4**). These results differ from those obtained by Nagata and coworkers (28), who injected a *Malat1* gapmer HDO at 50 mg/kg i.v. in rodents and found a robust knock-down activity in brain in cortex, brain stem and cerebellum in addition to the spinal cord. This discrepancy could be due to the different mechanism responsible for activity of *Malat1 a*nd *SMN2* HDOs, i.e., RNase-H competent versus splice-switching, or to the respective expression levels of *Malat1* and *SMN2* in brain. It could be that the dose of *Malat1* HDO delivered in brain after a single 50 mg/kg i.v. injection was sufficient to attain knock-down through the catalytic

RNAse-H mechanism, whereas correction of *SMN2* pre-mRNA splicing required a higher amount not reached at the dose we investigated (40 mg/kg). An alternative explanation may be the difference in length and/or in ribose modifications between our HDO construct (25-nt) and the published *Malat1* HDO construct (16-nt), which may have impacted tissue biodistribution and notably uptake in brain regions.

Further investigations are required to gain a better understanding of these parameters, and to establish comprehensive structure-activity relationship (SAR). In particular, different HDO lengths, delivery routes, and repeated dosing should be tested to build a comprehensive SAR and achieve optimal tissue activity and bioaccumulation. We also hypothesise that an SAR study would help identify optimal combinations for tolerability, as we observed injection-related side effects with **MOE**_HDO_ upon intravenous injection. Incidentally, Nagata et al (28) found that *Malat1* HDO subcutaneous injections were better tolerated than intravenous injections, and this information could be tested in future experiments with splice-switching HDOs. For instance, a further study could assess **MOE**_HDO_ at a lower dose in a repeated injection protocol, to establish whether splice correction in spinal cord and in muscle tissue can be enhanced. Finally, endocytosis and trafficking aspects were not addressed in our study. Further work could shed more light on **MOE**_HDO_ and **PMO**_HDO_ processing *in cellulo* and explore how these duplexes are cleaved or unwound once in tissues.

Notwithstanding the exploratory nature of our study, achieving splice correction in spinal cord is a key result in the context of SMA, as spinal cord hosts most motor neurons involved in the disease pathology (45). Another compelling finding was the significant activity achieved by **MOE**_HDO_ in peripheral tissues, e.g. skeletal and heart muscle (**Figure 5**) The data is of relevance for SMA, as splice correction in peripheral tissues is demonstrated to bring therapeutic benefits and be important for survival in animal models (29). Additionally, the findings pave the way for HDO studies in the context of other neuromuscular disorders. Further research should be conducted to revisit ON sequences that underperformed in the transition from *in vitro* to *in vivo* studies and could benefit from the HDO scaffold for enhanced tissular delivery. A natural extension of this work will be to assess HDOs carrying DMD ON drugs. Splice correction of *DMD* transcripts in skeletal and cardiac muscle is critical for therapy of DMD (46) and has been notoriously difficult to achieve with current therapies (9).

## MATERIAL AND METHODS

### Oligonucleotide synthesis

PMO oligonucleotides were purchased from Gene Tools LLC (Oregon, USA). 2’-*O*-MOE and 2’-*O*-methyl/RNA mixmer oligonucleotides were purchased from ATDBio Ltd (Oxford, United Kingdom). Oligonucleotides tested in cell culture and mouse experiments are listed in **Table 1**. Sterile-filtered, sodium salt forms of oligonucleotides were used for animal injections. PCR and RT-qPCR probes and primers were purchased from Integrated DNA Technologies (London, United Kingdom) and are listed in **Table S1**.

**Table 1.**
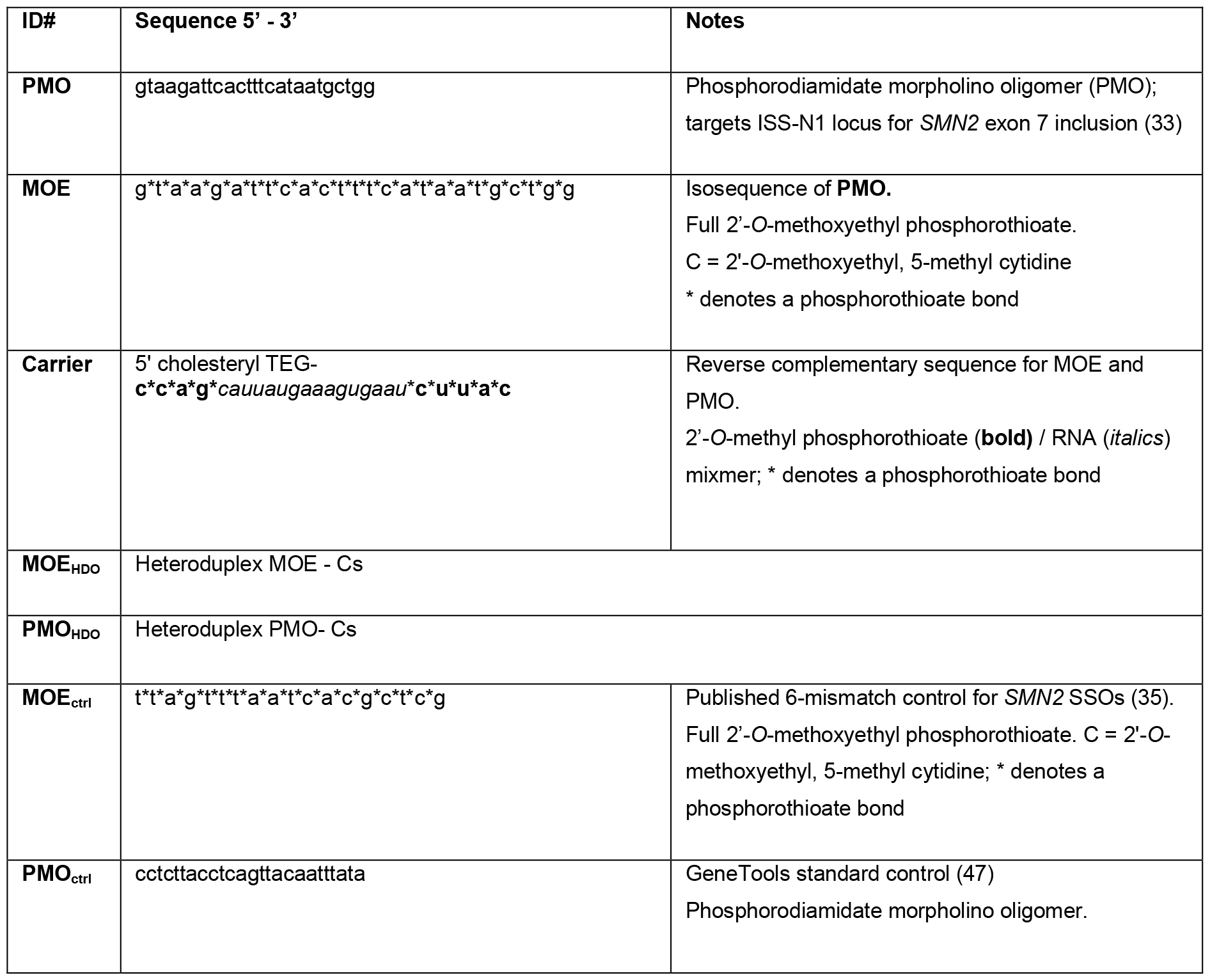
SMN2-targeting and control oligonucleotide sequences used for cell culture or animal experiments.

### Generation of heteroduplex oligonucleotides

Oligonucleotides were redissolved in RNase-free water and complementary strands mixed together at a final concentration of 100 µM. Annealing was performed on a thermocycler by heating at 95 C for 5 min and gradual cooling down to 25 C, with a cooldown rate of 0.1° C/second. Heteroduplex formation was verified by running annealed samples on 15% tris-borate-EDTA (TBE) gels for 1 h at 150 V followed by staining with SYBR Gold dye (1/10’000 dilution in water) for 10 minutes. By convention and in all experiments, concentrations of duplex solutions are expressed as concentrations of active strand within the duplex, so that comparisons between single-stranded and HDO oligonucleotides were made with equimolar amounts of active strand.

### Cell culture

#### General cell culture information

SMA Type II patient fibroblasts (Coriell Institute GM03813) were grown in DMEM-Glutamax (Gibco 31966-021) supplemented with 10 % fetal bovine serum (Gibco A5256701) and 1 % Antibiotic/Antimycotic solution (Merck A5955). Cells were kept in culture at 37°C, 5% CO_2_ and passaged using TrypLE Express (Gibco 12604013) twice weekly.

#### Testing of MOE oligonucleotides in SMA fibroblasts

Cells were plated at 75’000 cells/well in 500 µl medium in a 48-well plate. Oligonucleotide treatments were conducted 1.5 h after plating by transfection with Lipofectamine 2000 as per manufacturer’s instructions.

Oligonucleotides were pre-diluted in 25 µl OptiMEM medium immediately prior to mixing with 24 µl OptiMEM containing 1 µl Lipofectamine 2000. Solutions were incubated for 25 min before addition in cell culture medium. The total volume of oligonucleotide treatment solutions was 50 µl per well.

#### Testing of PMO oligonucleotides in SMA fibroblasts

Cells were plated at 50’000 cells/well in 500 µl medium in a 24-well plate. Cells were allowed to re-attach overnight before transfection with Lipofectamine 2000. Oligonucleotides were pre-diluted in 50 µl OptiMEM medium immediately prior to mixing with 49 µl OptiMEM containing 1 µl Lipofectamine 2000. Solutions were incubated for 25 min and added into cell culture wells. The total volume of oligonucleotide treatment solutions was 100 µl per well.

#### RNA extraction

Cells were washed one time with 500 µl phosphate-buffered saline (PBS) and pelleted. RNA extraction was carried out using Qiagen RNEasy kit according to manufacturer’s instructions and including the optional DNAse I treatment step. cDNA was generated using PrimeScript RT kit (Takara RR037A) with 100-500 ng RNA as input. cDNA samples were diluted 1/5 (v/v) in RNase-free water prior to use in real-time PCR.

#### Cell viability assay

Cell viability assays were conducted in 96 well-plate format using Promega’s RealTime-Glo MT Cell Viability Assay (Promega G9712) according to manufacturer’s instructions. Cells were plated at 10’000 cells/well in 200 µl medium in a 96-well plate and oligonucleotides delivered by Lipofectamine 2000 as in activity studies. Viability was measured 24 h post treatment in a microplate reader.

### Animal experiments

#### Animal strain and handling

The *hSMN2* transgenic mouse line (FVB.Cg-Smn1tm1HungTg(SMN2)2Hung/J) was maintained and treated at the Biomedical Sciences Unit, University of Oxford according to standard animal practices authorized by the UK Home Office (Animal Scientific Procedures Act, 1986). The mice were kept under a 12 h on / 12 h off light cycle and given free access to standard chow and purified water. These mice carry the wild-type mouse *Smn1* gene and four copies of the human *SMN2* gene.

#### Animal injections

Oligonucleotides for animal experiments were sterile-filtered over 0.2 µm acetate cellulose spin filters. Dilution with sterile concentrated saline solution to a final concentration of 0.9% saline was performed immediately prior to injection. Oligonucleotides were injected intravenously in the tail vein into 6–12-week-old mice at a dose of 40 mg/kg. Animals were monitored post injection for side effects and recovery time (if any). The human compassionate endpoint was incomplete recovery at 120 min post-injection. Seven days post administration, animals were culled via rising CO_2_ and decapitation. Cerebellum, cortex, brain stem, spinal cord, liver, heart, kidney, and tibialis anterior, gastrocnemius and quadriceps muscle tissues were collected. For spinal cord, the whole spinal column was excised from L5 to C2, followed by removal of muscle and surrounding soft tissue. The spinal column was then cut between C7 and T1 to liberate the cervical section, and between T13 and L1 to separate the lumbar section from the thoracic section. The vertebrae were then gently separated to expose the spinal cord for each of the sections. All tissues were snap frozen on dry ice and stored at –70°C for further analysis. Each dosage group had *n* = 4-5 mice per group.

#### RNA extraction from tissues

For all tissues bar skeletal muscle, tissue homogenization was carried out by bead beating and immediately followed by RNA extraction using the Maxwell® RSC simplyRNA Tissue Kit (Promega AS1340) as per manufacturer’s instructions. cDNA was generated using the High-Capacity cDNA Reverse Transcription Kit (Applied Biosystems 4368813) on a 500 ng or 1 µg scale total RNA, as determined by NanoDrop Microvolume Spectrophotometer (ThermoFisher Scientific). For skeletal muscle, tissue homogenization was carried out by bead beating and immediately followed by RNA extraction using the RNeasy Fibrous Tissue Mini Kit (Qiagen 74704), according to manufacturer’s protocol. Initial dissociation was conducted RLT buffer (Qiagen) supplemented with dithiothreitol, as per the manufacturer’s protocol. cDNA was generated using the Takara PrimeScript RT Reagent Kit and 500 ng total RNA input. cDNA samples were diluted 1/5 (v/v) in RNase-free water prior to use in semi-quantitative PCR or RT-qPCR.

#### Clinical Chemistry

Serum samples were analysed for total protein content, bilirubin concentration, and activity of alanine transaminase (ALT), alkaline phosphatase (ALP), and aspartate transaminase (AST) enzymes at the Clinical Chemistry platform of the Mary Lyon Centre (MRC Harwell, United Kingdom).

### Molecular biology assays

#### Mouse serum stability assay

Serum from BALB/C male mice, unfiltered, and complement preserved, was purchased from BioIVT. Ten nmoles of oligonucleotides were spiked into 300 µl serum and incubated for 24 h at 37°C with gentle shaking. References for t = 0 min timepoints were generated by taking an aliquot immediately upon spiking oligonucleotides into serum. Aliquots of 1.5 nmole were taken after 30-, 90-, or 180-min incubation and at t = 24 h. Aliquots were immediately mixed 1:1 (v/v) with RIPA buffer and 80 µg of proteinase K were added. Aliquots were digested for 30 min at 55°C prior to proteinase K inactivation for 5 min at 95°C and were frozen until analysis by PAGE on a 15% TBE gel. References for t = 0 min timepoints were generated by taking an aliquot immediately upon spiking oligonucleotides into serum and processing as described above. Samples were mixed 1:1 with 2x RNA dye (NEB B0363S), loaded on TBE gels, and run for 10 min at 85 V followed by 1 h at 140 V. Gels were stained for 10 min with SYBR Gold dye prior to imaging.

#### Quantification of full-length and total SMN2 transcripts by RT-qPCR

RT-qPCR reactions were prepared with the TaqMan™ Gene Expression Master Mix (Applied Biosystems # 4369016) and run and analysed on the StepOnePlus™ real-time PCR system (Applied Biosystems). Full-length *SMN2*, total *SMN2*, housekeeping *PolJ* and *ACTB* amplicons were amplified using TaqMan primers described in **Table S1**. *PolJ* was used as reference control in murine RNA samples and *ACTB* or total *SMN2* in human cell line samples. Each reaction was performed in duplicate. Fold changes were calculated using the ΔΔCt method. In human cell line samples, untreated cells were used as calibrators; in mouse experiments, saline-injected mice were used as calibrators.

#### Relative quantification of SMN2 full-length transcripts by semi-quantitative PCR

For semi-quantitative RT-PCR; PCR primers for simultaneous detection of *SMN2-Δ7* and full-length (FL) transcripts were used: Forward (FW) 5′-AAGTGAGAACTCCAGGTCTCCTG −3′, Reverse (RV) 5′-GTGGTGTCATTTAGTGCTGCTC −3′. A PCR mix containing 1 μl FW primer (10 μM stock), 1 μl RV primer (10 μM stock), 0.125 μl dNTPs mix (100 mM, Promega), 0.25 μl 5× Q5 Polymerase Buffer (NEB #M0491), 0.25 μl Q5 DNA Polymerase (NEB #M0491) and 15.75 μl ultrapure water was prepared and mixed with 1 μl of cDNA solution. PCR program: 98°C, 30 s; 34 cycles (98°C, 10 s – 69°C, 20s – 72°C, 2 s); 72°C, 2 min. PCR products were resolved by agarose gel electrophoresis in TAE (Tris-acetate-EDTA) buffer, using 2% agarose gels. Band intensity was quantified using ImageJ software with background subtraction. Ratios were calculated as: intensity of the full length band/(intensities of full length + Δ7 SMN2 bands)

#### Statistical *Analysis*

Statistical significance was determined with a minimum of *n* = 3 biological replicates using GraphPad Prism 9 software and one-way ANOVA tests with Dunnett’s correction. Statistical comparisons were made to saline control group (mouse experiments) or to untreated cells (SMA Type II fibroblast experiments). In cell culture experiments, data from independent replicates were combined with the value of the control group returned to 1 in each case. P values < 0.05 were considered to be statistically significant. Statistical tests and error bars are detailed in figure legends.

## DATA AVAILABILITY STATEMENT

The data underlying this article are available in the article and in its online supplementary material. Further raw data underlying this article will be shared on reasonable request to the corresponding authors.

## ACKNOWLEDGEMENTS

This work was supported by the Swiss National Science Foundation [P2EZP3_ 199973, to F.H.]; and UKRI Medical Research Council [MR/N024850/1 to M.W.]. The authors would like to thank the Animal Facility staff for their assistance with the animals.

## AUTHOR CONTRIBUTIONS

F.H. and M.W. conceived the project; F.H. performed cell culture experiments; N.A. conducted animal experiments and tissue/serum collection; N.A. and F.H processed animal tissues and carried out downstream molecular biology assays; F.H. analysed and curated data; F.H., N.A. and M.W. wrote the manuscript.

## SUPPLEMENTARY DATA

Supplementary Data are available for this manuscript.

