## Supplementary information for "Systemic delivery of a splice-switching oligonucleotide heteroduplex corrects splicing in central nervous system and muscle in spinal muscular atrophy mice"

**Table S1.** *Primers and probes used for TaqMan qPCR*. *FL SMN2* = full-length *SMN2* transcripts; Z = Zen™ modification, Fam = 6 Carboxyfluorescein, Hex = Hexachlorofluorescein, BFQ = 3 Iowa Black™ FQ .

| | **Sequence 5'-3'** |
|---|---|
| *FL SMN2* reverse primer | 5'-TCGTTTCTTTAGTGGTGTCATTTA-3' |
| *FL SMN2* forward primer | 5'-TATCATACTGGCTATTATATGGGTTTT-3' |
| *FL SMN2* probe | 5'-FAM-AAGGAGAAAZTGCTGGCATAGAGCAGC-BFQ-3' |
| *Total SMN2* reverse primer | 5'-TCAGTGCTGTATCATCCCAAATG-3' |
| *Total SMN2* forward primer | 5'-CAGGAGGATTCCGTGCTGTT-3' |
| *Total SMN2* probe[1] | 5'-FAM-CGGCACAGGZCCAGAGCGATG–BFQ-3' |
| *PolJ* (mouse) forward primer | 5′-TTCGAGTCGTTCTTGCTCTTC-3′ |
| *PolJ* (mouse) reverse primer | 5′-GTGGTCTTCTTTGTTGATGGTG-3′ |
| *PolJ* (mouse) probe | 5′-HEX-AAGCAGGCGZTTGGGAACCTTAGT-BFQ-3′ |
| *ACTB* (human) forward primer | 5'-ACAGAGCCTCGCCTTTG-3' |
| *ACTB* (human) reverse primer | 5'-CCTTGCACATGCCGGAG-3' |
| *ACTB* (human) probe | 5′-HEX-TCATCCATGZGTGAGCTGGCGG-BFQ-3′ |

[1] *Total SMN2 assay*. In transgenic DVBC mice, the assay does not recognise mouse *SMN* and is specific for the human *SMN2* transgene. In human patient fibroblasts, the assay detects all *SMN2* transcripts and can possibly detect some residual *SMN1* transcripts.

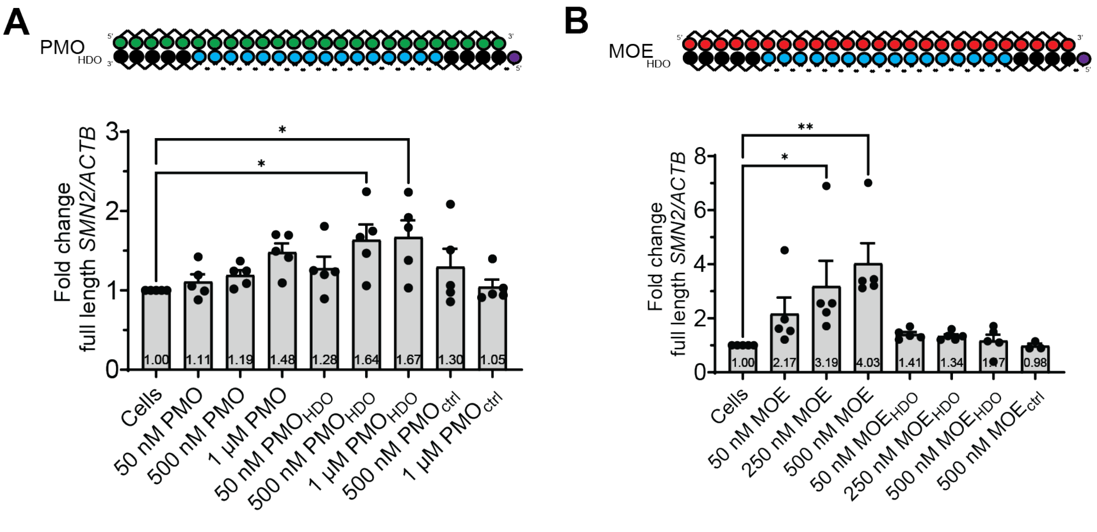

**Figure S1**. *In vitro activity of SMN2 splice-switching HDOs.*
Oligonucleotide activity was assessed in SMA Type II fibroblast cells after 24h treatment at indicated concentrations, by means of RT-qPCR assays detecting full-length *SMN2* and *ACTB* transcripts. Expression levels of full length SMN2 were normalised to total *SMN2* transcripts. **A** Cells were treated with increasing concentrations of **PMO**, **PMO**$_{HDO}$ or **PMO**$_{ctrl.}$ (n) = 5 biological replicates; data expressed as mean value (SEM: standard error of the mean); statistical analysis was conducted by comparison to the untreated group with one-way Anova (Dunnett's correction). **B** Cells were treated with increasing concentrations of **MOE**, **MOE**$_{HDO}$ or **MOE**$_{ctrl}$ (n) = 3-5 biological replicates; data expressed as mean value (SEM); statistical analysis was conducted by comparison to the untreated group with one-way Anova (Dunnett's correction).

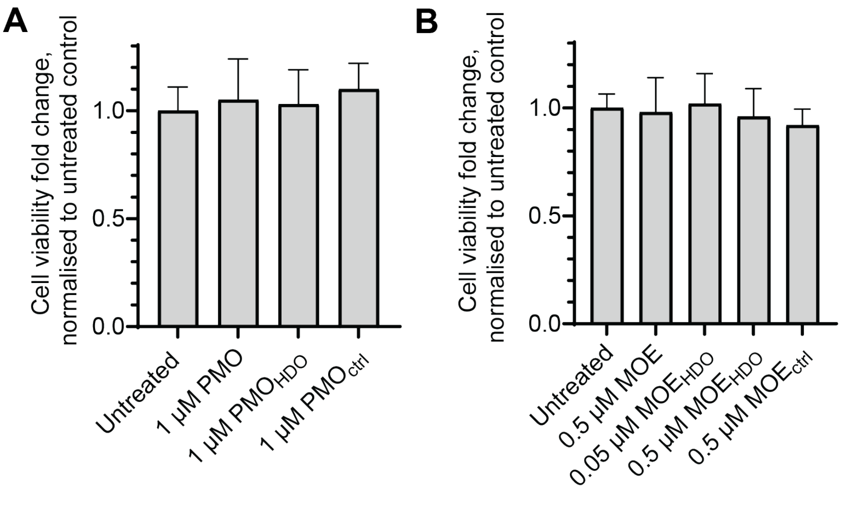

**Figure S2.** *Viability of SMA fibroblasts after oligonucleotide treatment.*
Fibroblasts were treated with **A:** 1 µM PMO, PMO$_{HDO}$ or PMO$_{ctrl}$, or **B:** 0.5 µM MOE, MOE$_{HDO}$ or MOE$_{ctrl}$, as detailed in methods section. Viability was assayed 24 h post treatment with Promega's RealTime-Glo MT Cell Viability Assay and luminescence measurement on a ClarioStar plate reader. Luminescence of treated samples was compared to untreated control. Data expressed as mean (SEM); *(n) = 3* replicates for Panel A, (*n*) = 4 replicates for Panel B.

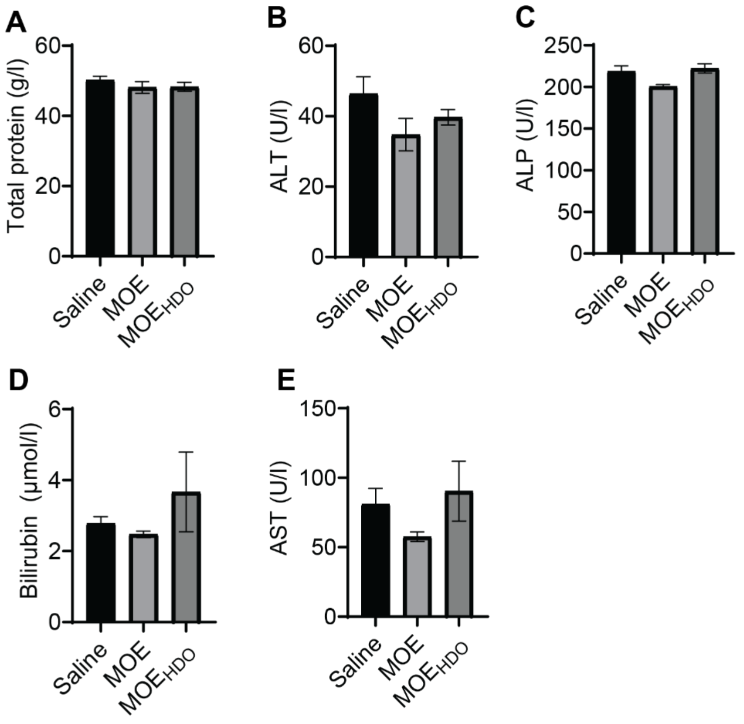

**Figure S3.** *Clinical chemistry biomarkers in SMA mice treated with a single dose of MOE or MOE$_{HDO}$ oligonucleotides.*

Animals were injected at 40 mg/kg intravenously with MOE or MOE$_{HDO}$ and culled seven days post injection. Serum was analysed for measurement of **A**: total protein content; **B**: activity of alanine transaminase (ALT), **C**: alkaline phosphatase (ALP), **D**: bilirubin concentration, and **E**: activity of aspartate transaminase (AST). Cohort size (*n*)= 3-5 per group; data expressed as mean value (SEM).

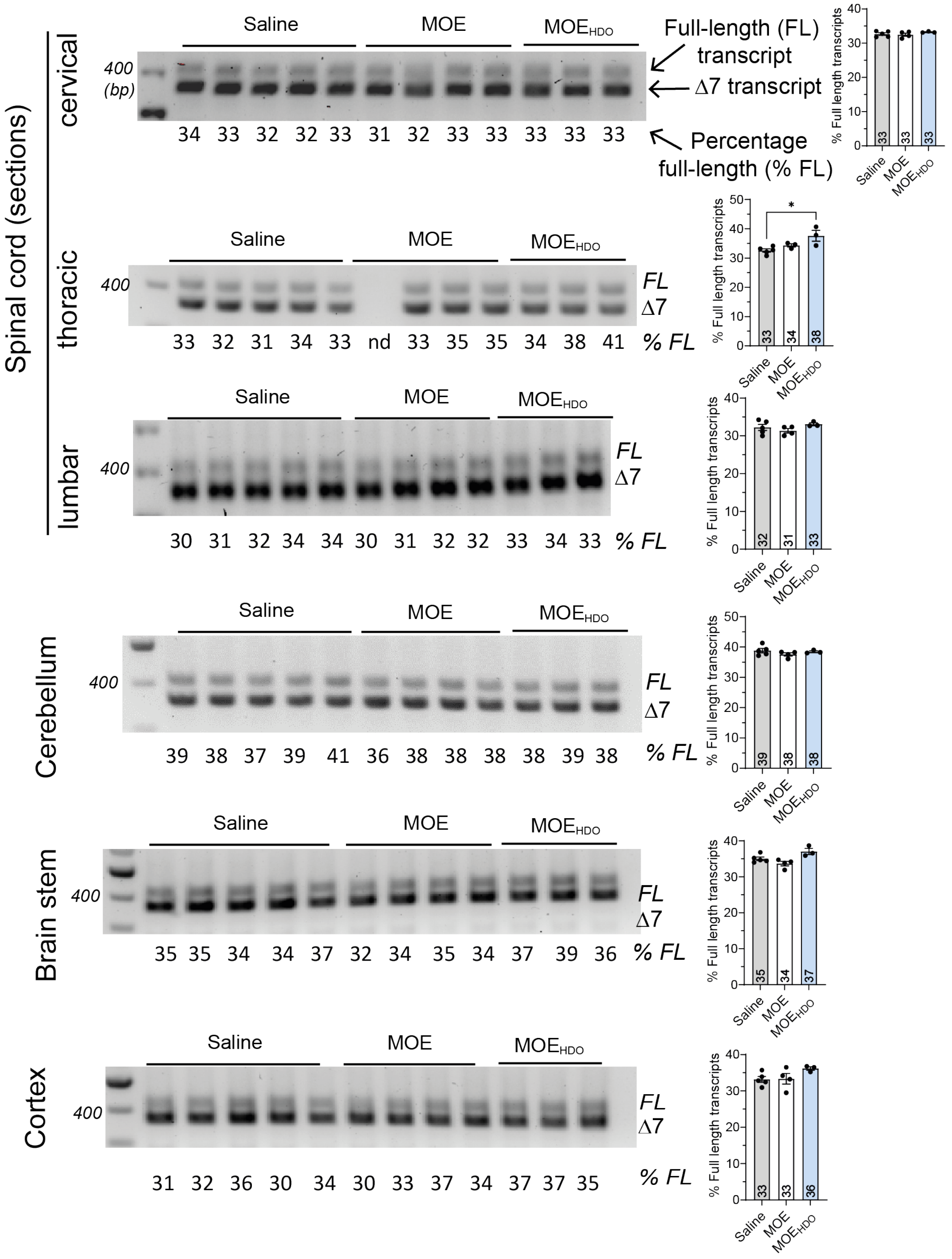

**Figure S4.** *The central nervous system activity of MOE and the corresponding heteroduplex $MOE_{HDO}$ in adult mice carrying the human SMN2 gene.*
Animals were injected at 40 mg/kg intravenously with MOE or $MOE_{HDO}$ and culled seven days post injection. Activity was assayed with a semi-quantitative PCR assay for simultaneous detection of correct and Δ7 (i.e., truncated) SMN2 transcripts. Left-hand-side: Full-length (FL) and truncated (Δ7) amplicons detected on agarose gel. Each lane represented an independent experimental animal. "nd" indicated a failed PCR reaction; no value was recorded. Right-hand-side: averaged % FL values for each tissue. Animal cohort size (*n*)= 3-5 per group; statistical significance was determined by comparison to the saline group with one-way ANOVA (Dunnett's correction); * = $p < 0.05$; data expressed as mean value (SEM).

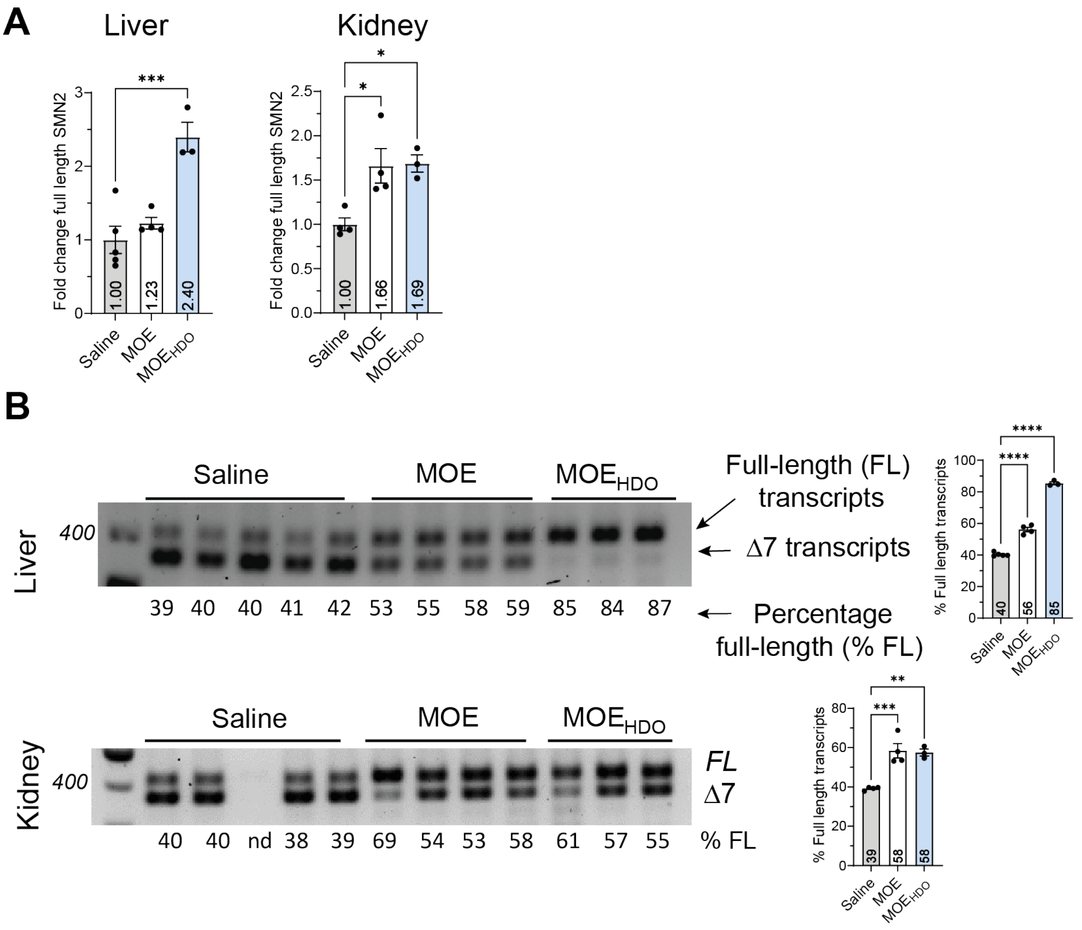

**Figure S5.** *The kidney and liver activity of the 25-nt splice-switching oligonucleotide MOE and the corresponding heteroduplex $MOE_{HDO}$ in adult mice carrying the human SMN2 gene***.** Animals were injected at 40 mg/kg intravenously with MOE or $MOE_{HDO}$ and culled seven days post injection. **A** Activity was assayed with a RT-qPCR assay for quantification of correct *SMN2* transcripts and normalization to the reference transcript *PolJ,* and **B** with a semi-quantitative PCR assay for simultaneous detection of full length and Δ7 *SMN2* transcripts. “nd” indicated a failed PCR reaction; no value was recorded. Right-hand-side: averaged % FL values for each tissue. Animal cohort size (*n*)= 3-5 per group; statistical significance was determined via one-way ANOVA with Dunnett’s correction; * = $p < 0.05$, ** = $p < 0.01$, *** = $p < 0.001$; **** = $p < 0.0001$.; data expressed as mean value (SEM).
